# Genome-scale CRISPR Screens Identify Host Factors that Promote Human Coronavirus Infection

**DOI:** 10.1101/2021.06.04.447090

**Authors:** Marco Grodzki, Andrew P. Bluhm, Moritz Schäfer, Abderrahmane Tagmount, Max Russo, Amin Sobh, Roya Rafiee, Chris D. Vulpe, Stephanie M. Karst, Michael H. Norris

## Abstract

The COVID-19 pandemic has resulted in 153 million infections and 3.2 million deaths as of May 2021. While effective vaccines are being administered globally, there is still a great need for antiviral therapies as potentially antigenically distinct SARS-CoV-2 variants continue to emerge across the globe. Viruses require host factors at every step in their life cycle, representing a rich pool of candidate targets for antiviral drug design. To identify host factors that promote SARS-CoV-2 infection with potential for broad-spectrum activity across the coronavirus family, we performed genome-scale CRISPR knockout screens in two cell lines (Vero E6 and HEK293T ectopically expressing ACE2) with SARS-CoV-2 and the common cold-causing human coronavirus OC43. While we identified multiple genes and functional pathways that have been previously reported to promote human coronavirus replication, we also identified a substantial number of novel genes and pathways. Of note, host factors involved in cell cycle regulation were enriched in our screens as were several key components of the programmed mRNA decay pathway. Finally, we identified novel candidate antiviral compounds targeting a number of factors revealed by our screens. Overall, our studies substantiate and expand the growing body of literature focused on understanding key human coronavirus-host cell interactions and exploit that knowledge for rational antiviral drug development.

**One Sentence Summary:** Genome-wide CRISPR screens identified host factors that promote human coronavirus infection, revealing novel antiviral drug targets.

## INTRODUCTION

The COVID-19 pandemic is arguably the most consequential infectious disease outbreak in modern times. The causative agent of COVID-19, SARS-CoV-2, spread quickly across the planet resulting in 153 million infections and 3.2 million deaths at the time of this writing. Multiple COVID-19 vaccines recently demonstrated high efficacy, received FDA approval, and are being administered to people across the globe. While the importance of this scientific achievement cannot be understated, there is still a great need for novel antiviral therapies for use in vulnerable immunocompromised individuals, in regions where vaccine access is limited, and in the event that antigenically distinct SARS-CoV-2 variants emerge. Moreover, considering that SARS-CoV-2 is the third novel human coronavirus (HCoV) to emerge and cause serious disease in the human population in the past two decades following SARS-CoV and MERS-CoV, potent and broad-spectrum antivirals will leave us better prepared to deal with future pandemics. Broad-spectrum antivirals could also reduce morbidity associated with common cold-causing HCoVs including OC43, NL63, 229E, and HKU1.

Antivirals segregate into two basic categories, virus-targeting and host-targeting, both of which require an understanding of the molecular mechanisms used by viruses to replicate in host cells. Coronaviruses replicate via a well-established series of molecular events *(1, 2).* Host factors are required at every step in this life cycle and represent candidate druggable targets (i.e. host-targeting antivirals) with the potential for broad-spectrum activity against multiple viruses within a given virus family and even across virus families *(3, 4).* Accordingly, we performed CRISPR-based genome-wide knockout screens for both SARS-CoV-2 and OC43 infections to identify host factors that promote HCoV replication. Considering the power of genome-wide screens in the identification of host factors required for viral replication and the enormous global impact of the ongoing COVID-19 pandemic, it is not surprising that other research groups also applied this approach to SARS-CoV-2. Six genome-wide CRISPR screens for the identification of host factors promoting SARS-CoV-2 replication are published *(5–10).* Despite the redundancy in overall approach, there was experimental variability across screens in the selection of cell lines and infection conditions. Together with the sheer magnitude of critical host-virus interactions required for successful viral infection, individual screens are likely to capture only a subset of these host factors. Consistent with this, while specific genes and pathways were identified across published studies, each study also provided unique findings which expand our understanding of host-HCoV interactions.

In this study, we report a global analysis of host-HCoV interactions gleaned from genome-wide screens performed for two HCoVs and in two different cell lines. We also performed a comprehensive comparative analysis of all published genome-wide SARS-CoV-2 screens to date. Multiple genes and functional pathways identified in our screens were previously reported to promote SARS-CoV-2 replication, validating the rigor of our approach and providing further support for the role of specific host factors. Yet we also identified a substantial number of novel genes and pathways not previously reported to promote HCoV replication. We validated the importance of a subset of genes identified in these screens in HCoV replication. Notably, several of the novel host factors identified in our study provide unique insight into SARS-CoV-2 replication processes that could be targeted with antiviral drugs. Host factors involved in cell cycle regulation were enriched in our screens and we show that compounds (abemaciclib, AZ1 protease inhibitor, harmine, nintedanib, and UC2288) targeting these host factors inhibit *in vitro* HCoV replication. We also identified multiple host factors involved in endocytosis and TBK1 that plays a key role in innate immune responses. Inhibitors of these processes/factors (promethazine and amlexanox, respectively) also displayed antiviral activity. Together, our study has provided significant insight into host-HCoV interactions and identified novel candidate antiviral compounds.

## RESULTS

### Genome-wide CRISPR screens in Vero E6 cells identify host factors required for HCoV infection

In order to identify host factors that promote HCoV infection, we performed genome-wide loss of function CRISPR screens for pathogenic SARS-CoV-2 and common cold-causing OC43 in two susceptible cell lines. Due to the highly cytopathic nature of HCoV infection in the Vero E6 cells derived from African Green Monkey (AGM; *Chlorocebus sabaeus*), we carried out genome-wide screens using a custom Vervet CRISPR knockout library (see supplemental methods) **(Fig. 1A)**. Vero E6 cells were transduced with the Vervet CRISPR library and infected with SARS-CoV-2 or OC43 at a multiplicity of infection (MOI) 0.01. We observed ~60% visible cytopathic effect (CPE) for SARS-CoV-2 and ~85% CPE for OC43. Resistant clones were expanded, reinfected with the corresponding virus at MOI 0.1, and re-expanded. Genomic DNA was extracted from surviving cells, sgRNAs amplified, and sequenced. We carried out MAGeCK analysis to identify genes targeted by significantly enriched sgRNAs which are labeled in the volcano plots in **Figs. 2A-B**. To facilitate the access and reusability of data sets generated from genome-scale CRISPR screens in HCoV-infected cells, we have also hosted a website (sarscrisprscreens.epi.ufl.edu) with the complete set of MAGeCK results for each of the screens described in this study and data from previously published screens reanalyzed herein *(3, 7–10).* The website was designed to facilitate integration of upcoming screens and we hope for contributions to drive this as a community project.

**Figure 1.**
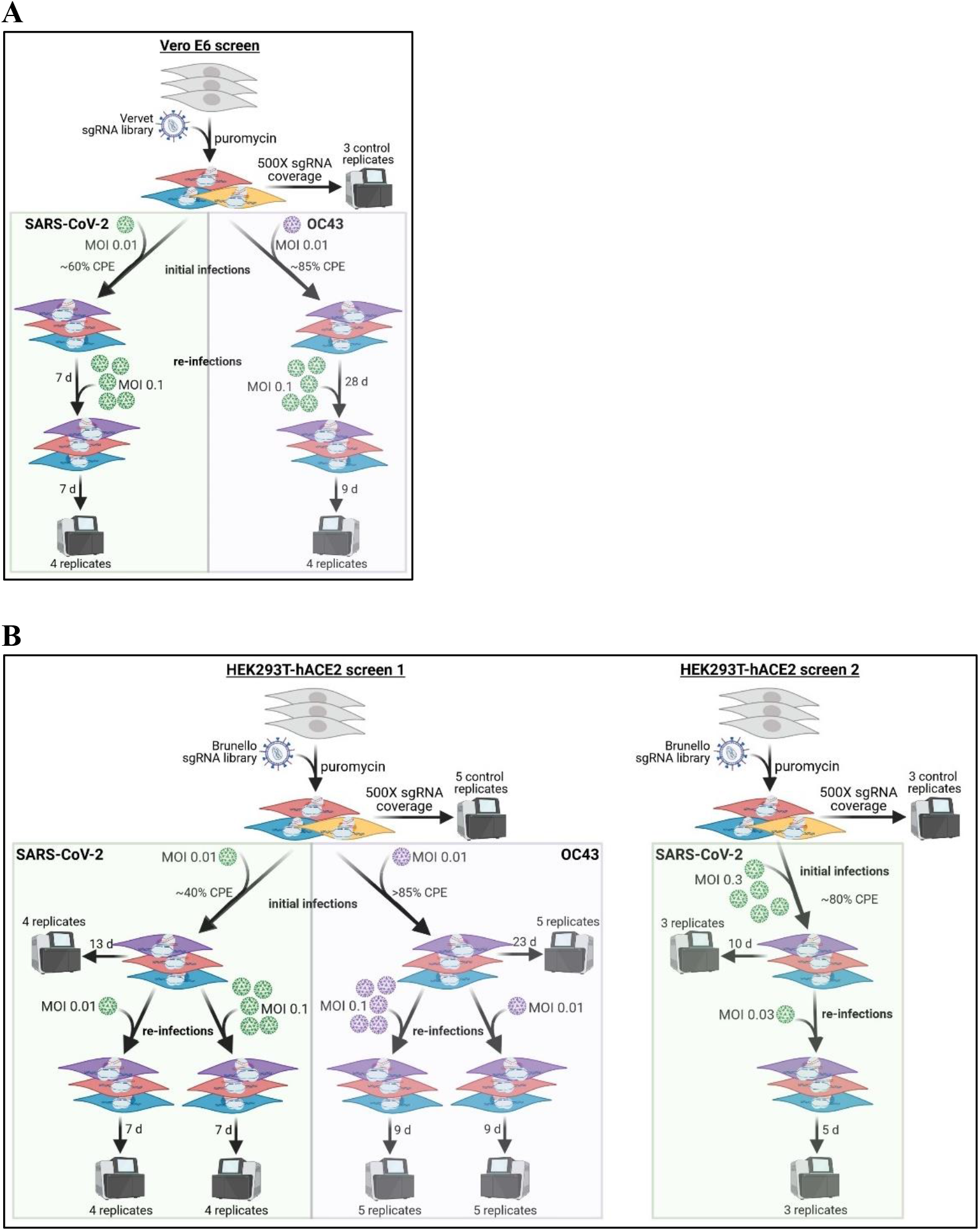
Experimental design for genome-scale CRISPR screens performed in this study. Details of these screens are provided in the methods. **A)** Vero E6 cells transduced with the newly generated Vervet sgRNA library were infected with SARS-CoV-2 or OC43 at MOI 0.01, resistant cells were expanded and re-infected at MOI 0.1 **B)** Two screens were performed in HEK293T-hACE2 cells transduced with the Brunello sgRNA library. In the first screen, cells were infected with SARS-CoV-2 or OC43 at MOI 0.01 and resistant cells were re-infected with either MOI 0.01 or MOI 0.1 of the corresponding virus. In the second screen, cells with infected with SARS-CoV-2 at MOI 0.3 and re-infected at MOI 0.03. In all cases, genomic DNA was extracted from multiple replicates of control cells, the initial infections, and re-infections for the purpose of sgRNA sequencing.

**Fig 2.**
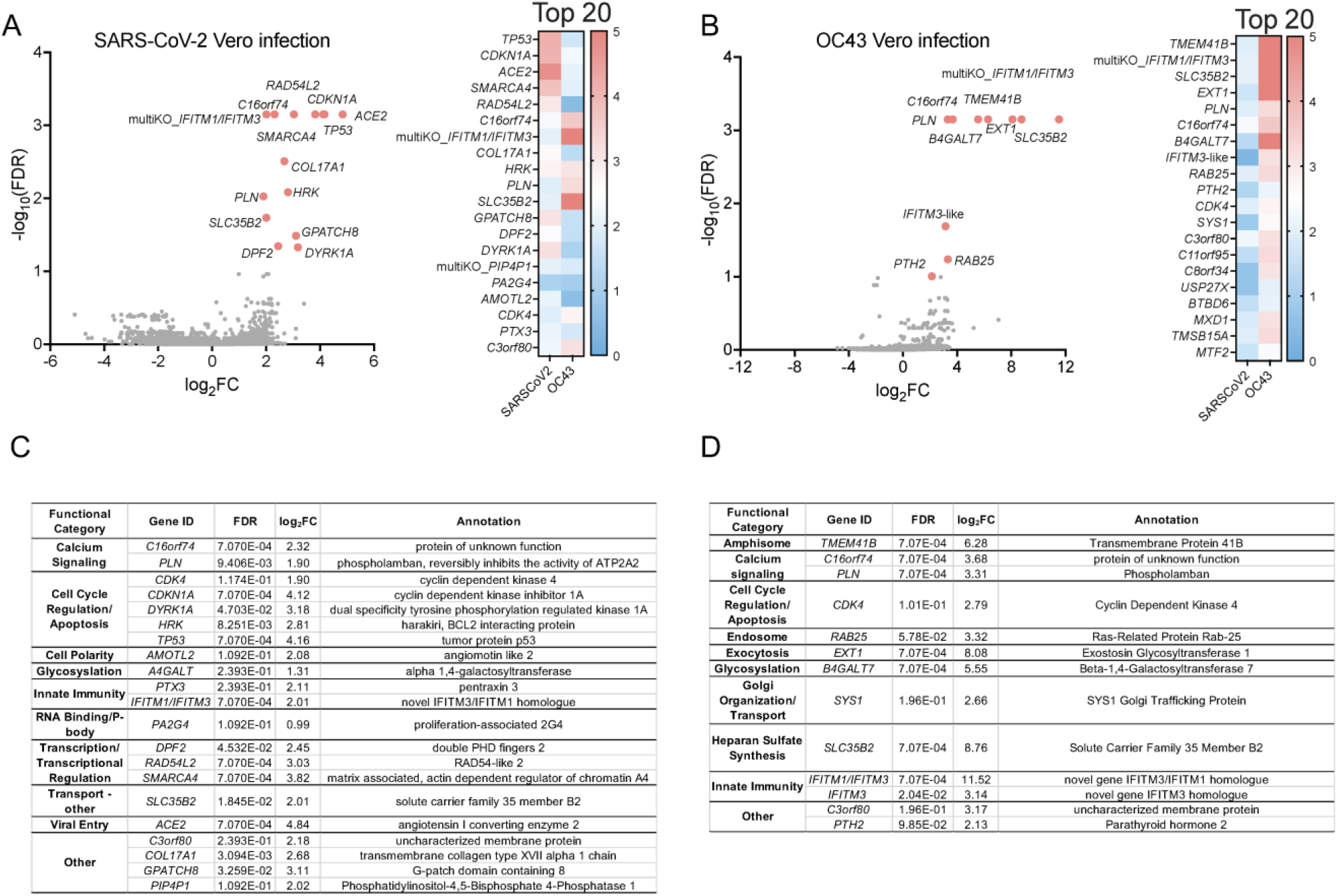
Identification of host factors that promote HCoV infection of Vero E6 cells. Vero E6 cells transduced with a *C. sabaeus*-specific sgRNA library were infected with SARS-CoV-2 or OC43 at MOI 0.01, resistant cells reinfected at MOI 0.01, and sgRNAs in resistant clones sequenced. MAGeCK analysis of multiple replicates compared to uninfected control library replicates yielded log_2_fold changes (log_2_FC) that were plotted on the x-axis. Negative log10 transformed false discovery rates (FDR) were plotted on the y-axis. Data are presented for SARS-CoV-2 **(A)** and OC43 **(B)** Vero E6 infections. The heat maps display the log_2_FC for the 20 top-scoring genes comparing results for SARS-CoV-2 and OC43 infections. Genes targeted by significantly enriched sgRNAs (FDR<0.25) were segregated into functional categories using PANTHER for SARS-CoV-2 **(C)** and OC43 **(D)** infections.

We identified multiple candidate host factors previously demonstrated to play a functional role in SARS-CoV-2 and OC43 infections. For example, *ACE2* was identified in the SARS-CoV-2 screen *(11).* Furthermore, *TMEM41B* was a top-scoring gene in the OC43 screen, supporting recent work by Schneider et al. demonstrating that *TMEM41B* is a pan-HCoV host factor *(7)*. We also identified interferon (IFN)-induced transmembrane (IFITM) proteins that have been reported to regulate HCoV infection *(12–14).* Genes targeted by significantly enriched sgRNAs (FDR<0.1) were next segregated into functional categories listed in tables in **Figs. 2C-2D**. Of note, *CDK4*, a master regulator of cell cycle, was identified as a key host factor for both viruses. Disruption of additional genes encoding regulators of cell cycle progression, including *CDK1NA*, *DYRK1A*, *HRK* and *P53*, similarly increased cellular resistance to SARS-CoV-2 infection.

During the completion of our studies, Wei et al. reported similar SARS-CoV-2 screens in Vero E6 cells performed with an independent sgRNA library based on an earlier *C. sabaeus* genome assembly *(8).* In order to compare our data sets to those of Wei et al., we downloaded raw data from their study and analyzed them using MAGeCK-VISPR *(8, 15).* There were 6 targeted genes identified in common between studies: *ACE2, DPF2, DYRK1A, RAD54L2, SMARCA4, and TP53*.

### Genome-wide CRISPR screens in HEK293T-hACE2 cells identify host factors required for HCoV infection

We similarly performed CRISPR screens in human HEK293T cells ectopically expressing the human ACE2 receptor (HEK293T-hACE2) transduced with the Brunello sgRNA library *(16)* **(Fig. 1B)**. Transduced cells were infected with SARS-CoV-2 or OC43 at MOI 0.01. SARS-CoV-2-infected cultures developed ~40% CPE and OC43-infected cultures developed >85% CPE. Resistant cell populations propagated to confluence were reinfected with the corresponding virus at either MOI 0.01 or MOI 0.1 and re-expanded. Genomic DNA was extracted and sgRNAs from both the initial and secondary infections were sequenced.

The genes targeted by the most highly enriched sgRNAs in each of the SARS-CoV-2 infections are indicated in **Figs. 3A-3C**. *EDC4*, a gene encoding a scaffold protein that functions in programmed mRNA decay, was the overall top-scoring gene. Interestingly, *XRN1* encodes another key player in this pathway and was also top-scoring. We further categorized the genes encoding candidate host factors (FDR<0.1) into functional categories depicted in heat maps in **Fig. 3D**. Consistent with other published screens, we identified multiple components of the endocytic pathway including *CCZ1*, *DNM2*, and *WASL*. Other functional categories in which multiple genes were identified include cell adhesion, cell cycle, integrator complex, lysosome, mTOR regulation, and ubiquitination/proteolysis. We carried out an independent SARS-CoV-2 screen using the higher MOI of 0.3 for initial infection which resulted in ~80% CPE, and MOI 0.03 for secondary infection. Genes targeted by the most significantly enriched sgRNAs in this study are presented in **Figs. 3E-3F** and are segregated into functional categories depicted in heat maps in **Fig. 3G**. *ACE2* was a top-scoring gene in this screen. Functional categories with multiple targeted genes include amphisome, autophagy, endosome, exocytosis, lysosome, peroxisome, transcription/transcriptional regulation, and ion transporters. *C18orf8, CCZ1, CDH2*, and *TMEM251* were identified in both the low- and high-MOI SARS-CoV-2 screens.

**Fig 3.**
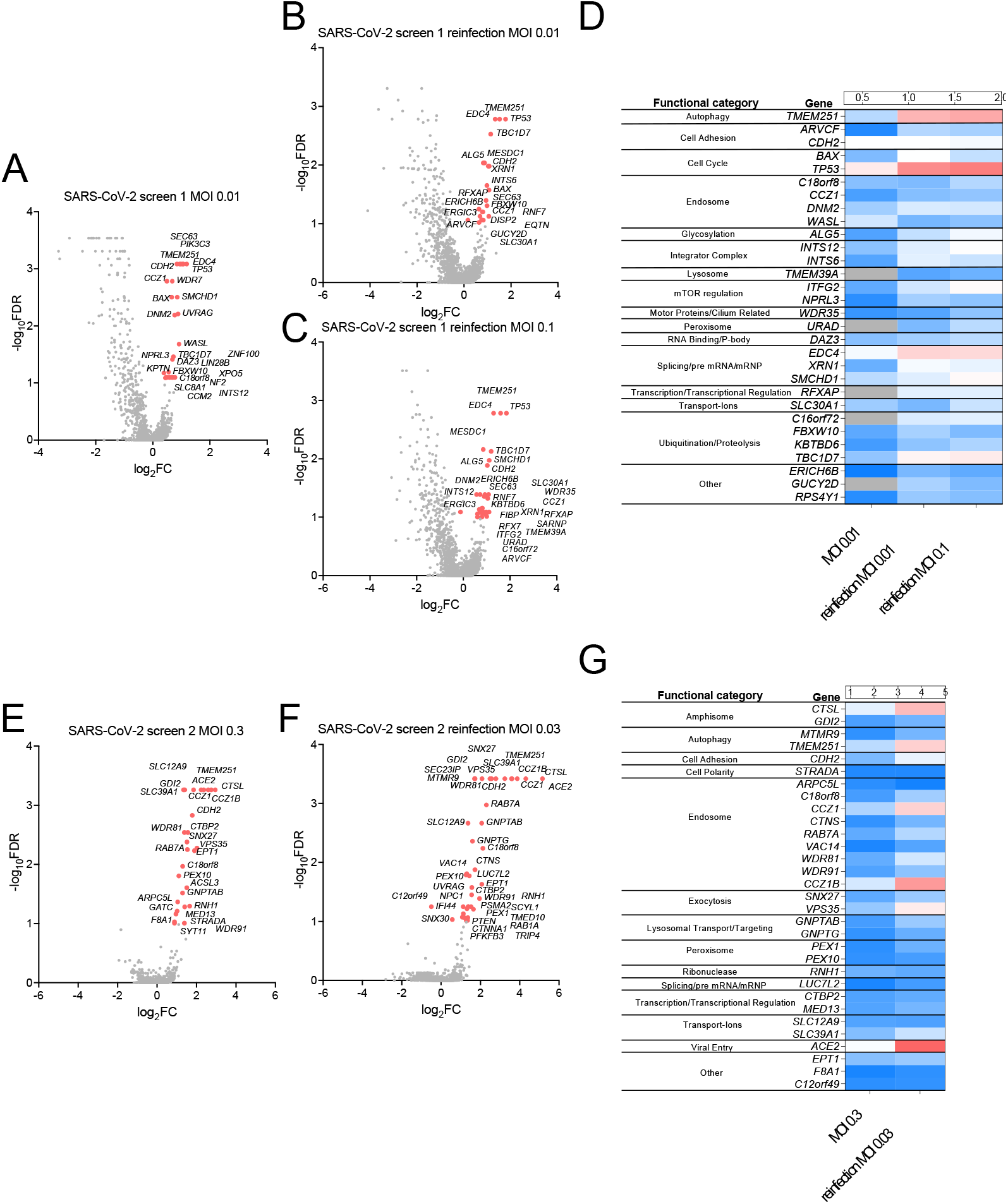
Identification of host factors that promote SARS-CoV-2 infection of HEK293T-hACE2 cells. HEK293T-hACE2 cells transduced with the Brunello sgRNA library were infected with SARS-CoV-2 at MOI 0.01 and sgRNAs in resistant clones sequenced. Resistant clones were reinfected with SARS-CoV-2 at MOI 0.01or MOI 0.1 and sgRNAs in resistant clones sequenced. For all three infections, MAGeCK analysis of multiple replicates compared to uninfected control library replicates yielded log_2_fold changes (log_2_FC) that were plotted on the x-axis. Negative log10 transformed FDR were plotted on the y-axis. Data are presented for the initial infection **(A)**, MOI 0.01 reinfection **(B)**, and MOI 0.1 reinfection **(C)**. **D)** The heat map displays the log_2_FC for top-scoring genes (FDR<0.1) across the three infections. The entire experiment was repeated at MOI 0.3, with sgRNAs sequenced from resistant clones in the initial infection **(E)** and reinfection **(F)**. **G)** The heat map displays the log_2_FC for top-scoring genes (FDR<0.25) across the two infections.

The genes targeted by the most highly enriched sgRNAs in the OC43 HEK293T-hACE2 screens are indicated in **Figs. 4A-4C** and segregated into functional categories in **Fig. 4D**. As expected, based on prior work, genes encoding IFITM proteins were identified as proviral factors for OC43 *(12)*. *TMEM41B* was a top-scoring gene along with the functionally related *VMP1*, as were *CCZ1, CCZ1B, SLC35B2*, and *WDR81* which have all been reported in other recent OC43 genome-wide screens *(6, 7).* When comparing the SARS-CoV-2 and OC43 HEK293T-hACE2 datasets, there were 6 genes in common targeted by significantly enriched sgRNAs (*C18orf8, CCZ1, CCZ1B, RAB7A, WDR81*, and *WDR91*). Notably, all of the corresponding gene products function in vesicle-mediated transport.

**Fig 4.**
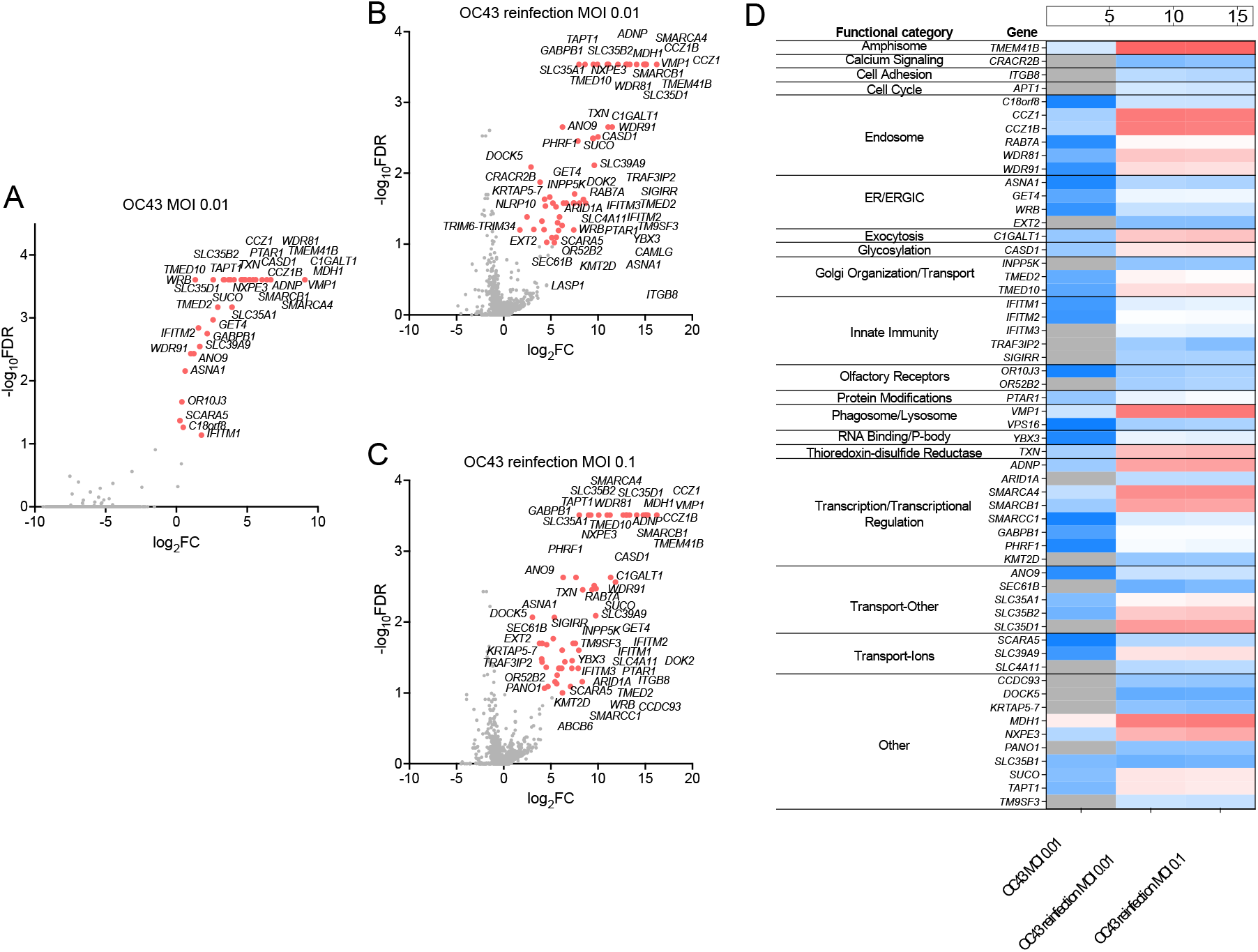
Identification of host factors that promote OC43 infection of HEK293T-hACE2 cells. HEK293T-hACE2 cells transduced with the Brunello sgRNA library were infected with OC43 at MOI 0.01 and sgRNAs in resistant clones sequenced. Resistant clones were reinfected with OC43 at MOI 0.01or MOI 0.1 and sgRNAs in resistant clones sequenced. For all three infections, MAGeCK analysis of multiple replicates compared to uninfected control library replicates yielded log_2_fold changes (log_2_FC) that were plotted on the x-axis. Negative log10 transformed FDR were plotted on the y-axis. Data are presented for the initial infection **(A)**, MOI 0.01 reinfection **(B)**, and MOI 0.1 reinfection **(C)**. **D)** The heat map displays the log_2_FC for top-scoring genes (FDR<0.25) across the three infections.

During the completion of our studies, similar SARS-CoV-2 screens in human Huh-7.5 *(6, 7, 10)* and A549 *(5, 9)* cells were published. In order to compare our data sets to those of other groups, we downloaded raw data from four published studies *(6, 7, 9, 10)*, analyzed them using a common analysis framework (MAGeCK) and stringency (FDR<0.25) and compared the results to our data sets. Using this stringency, no genes were identified in all five studies, 1 gene was identified in four studies (*ACE2*), 6 genes were identified in three studies (*VPS35, CTSL, DNM2, CCZIB, TMEM106B*, and *VAC14*), and 25 genes were identified in two studies (*ALG5, ARVCF, ATP6V1A, ATP6V1G1, B3GAT3, CNOT4, EPT1, EXOC2, EXT1, EXTL3, GDI2, LUC7L2, MBTPS2, PIK3C3, RAB7A, RNH1, SCAF4, SCAP, SLC30A1, SLC33A1, SNX27, TMEM41B, TMEM251, WDR81*, and *WDR91*) **(Fig. 5A-5B)**. It should be noted that these genes were top-scoring across studies performed in different human cell lines, suggesting they are broadly important in SARS-CoV-2 replication. Shared pathways include vesicle-mediated transport (*CCZ1B, DNM2, EXOC2, GDI2, PIK3C3, RAB7A, SNX27, VAC14, VPS35, WDR81, WDR91*), vacuolar ATPases important in organelle acidification (*ATP6V1A, ATP6V1E, ATP6V1G1*), and heparan sulfate biosynthesis genes (*EXT1, EXTL3, B3GAT3*). We identified 53 genes targeted by enriched sgRNAs in our study that were not identified in published studies **(Fig. 5C)**, including *EDC4* and *XRN1*.

**Fig 5.**
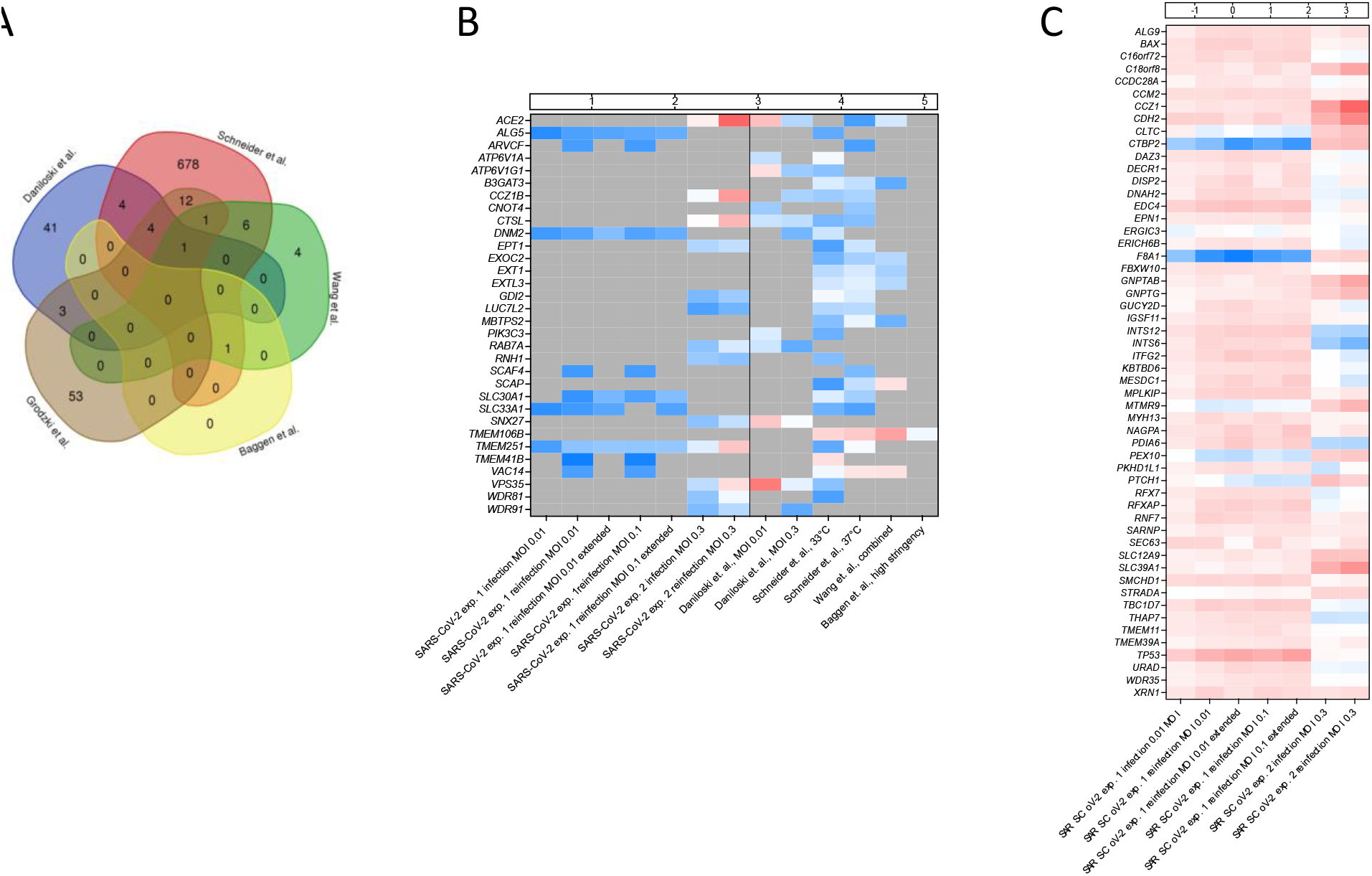
Comparison of multiple CRISPR screens identifying host factors promoting SARS-CoV-2 infection of human cell lines. **A)** Data from four recently published CRISPR screens for SARS-CoV-2 in various human cell lines were reanalyzed and compared to our data to identify common top-scoring genes (FDR<0.25). Using this criterion, there were 74 genes identified in our study, 53 in Daniloski et al., 707 in Schneider et al., 13 in Wang et al., and 1 in Baggen et al. No common genes were identified in all studies, 1 gene was identified in four studies, 6 genes were identified in three studies, and 25 genes were identified in two studies. 53 genes were uniquely identified in our study as significant. **B)** The heat map displays the log_2_FC for the 32 genes found in common across two or more of the published studies with FDR<0.25. **C)** A heat map displaying the log_2_FC for the 53 genes uniquely identified as significant in our studies compared to their observed log_2_FC across the other published studies.

### Validation of a subset of gene candidates that promote human coronavirus replication

To confirm that unique genes identified in our screens promote HCoV replication, HEK293T-hACE2 cells were engineered to stably express gene-specific shRNAs targeting *CCZ1* or *EDC4. CTSL* knockdown was tested as a positive control for SARS-CoV-2 *(17).* Efficiency of gene knockdown assessed by western blotting was robust for all three genes **(Fig. 6A)**. Knockdown cells were then infected with SARS-CoV-2 or OC43 and viral genome copy number determined at 2 days post-infection (dpi). All three genes were required for optimal SARS-CoV-2 infection while *CCZ1* and *EDC4*, but not *CTSL*, promoted OC43 infection **(Fig. 6B)**.

**Fig 6.**
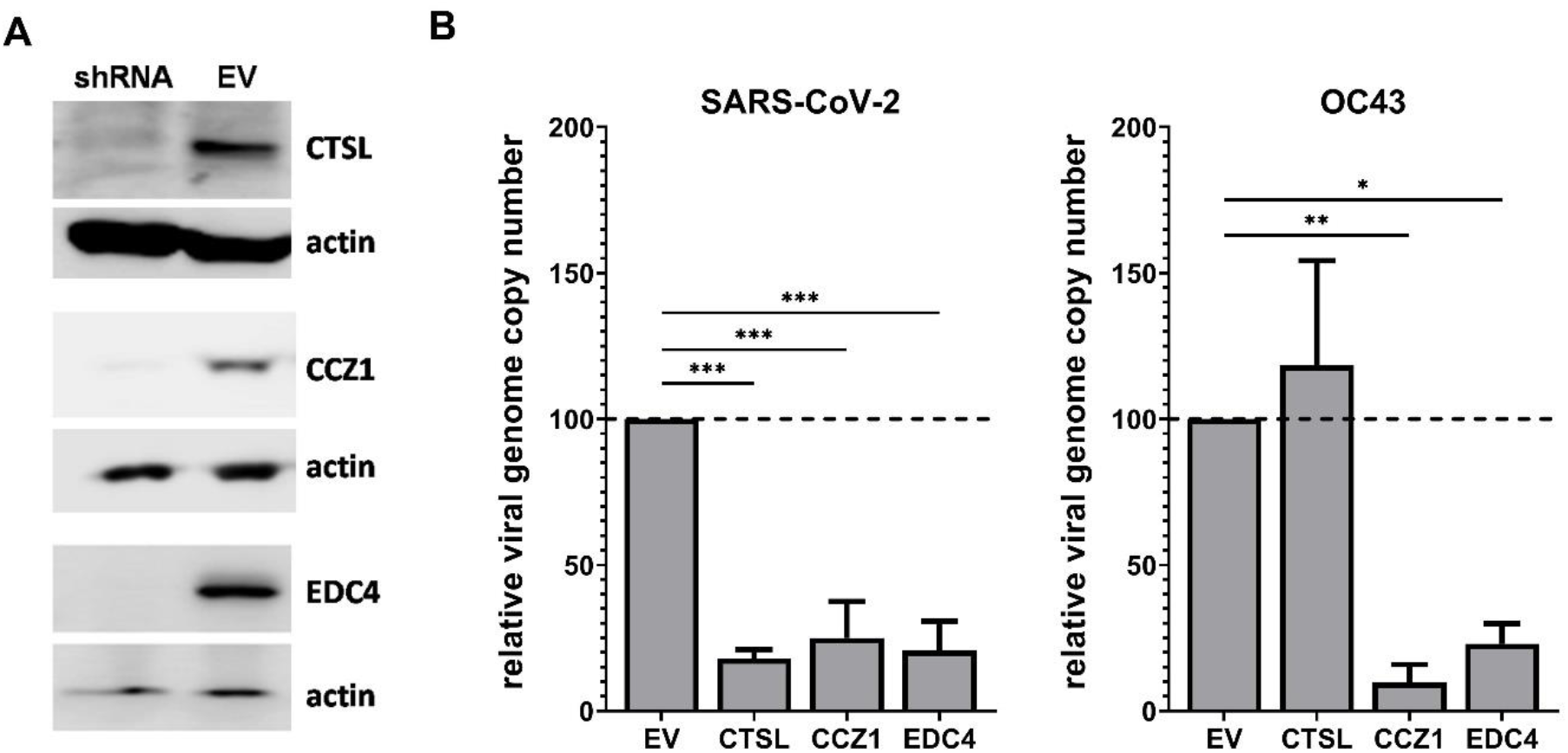
Confirmation of host factor involvement by targeted shRNA knockdown. Lentivirus-packaged shRNA clones directed to *CTSL*, *CCZ1*, and *EDC4* were transduced into HEK293T-hACE2 cells and selected with puromycin. **A)** Gene knockdown was assessed using western blotting with antibodies directed to CTSL, CCZ1, and EDC4 in cells transduced with a gene-specific shRNA or empty vector control (EV). Actin expression served as a loading control. **B)** Triplicate wells of knockdown cells were infected with SARS-CoV-2 or OC43 at MOI 0.01. At 2 dpi, viral genome copy numbers were determined by RT-qPCR and normalized to *GAPDH* levels as a housekeeping control. The data are reported as the relative normalized viral genome copy number in shRNA-expressing cells compared to the EV control (*n* = 3 experiments). Error bars denote standard errors of mean and *P* values were determined using one-way ANOVA (\**P* < 0.05, \*\**P* < 0.01, \*\*\**P* < 0.001).

### CRISPR screening reveals novel antiviral drugs displaying *in vitro* efficacy

We next determined whether gene products and pathways identified in our screens could be targeted with commercially available inhibitors to block HCoV infection. Numerous genes involved in cell cycle regulation were identified in our screens. The following inhibitors targeting this class of host factors were tested: abemaciclib (ABE; Cdk4 inhibitor), UC2288 (UC2; CDKN1A/p21 inhibitor), harmine (HAR) and INDY (Dyrk1A inhibitors), AZ1 (Usp25/28 inhibitor), olaparib (OPB; ARID1A inhibitor), and nintedanib (NIN; FGFR1/2/3 inhibitor). Host factors involved in endocytosis have been widely reported to regulate HCoV replication and were identified in our and others’ CRISPR screens *(18)* so we also tested several drugs targeting this process including CID1067700 (CID; Rab7a inhibitor), chlorpromazine (CPZ; Wdr81 inhibitor), and promethazine (PMZ; Wdr81 inhibitor). Finally, we tested amlexanox (AMX) which inhibits TANK binding kinase 1 (TBK1) and its adaptor protein TBK-binding protein 1 (TBKBP1) which has been reported to variously regulate Rab7a activity *(19)* or induction of IFN response genes *(20).* The heat map in **Fig. 7A** shows the fold-enrichment of sgRNAs targeting the genes of interest across the screens performed in this study.

**Fig 7.**
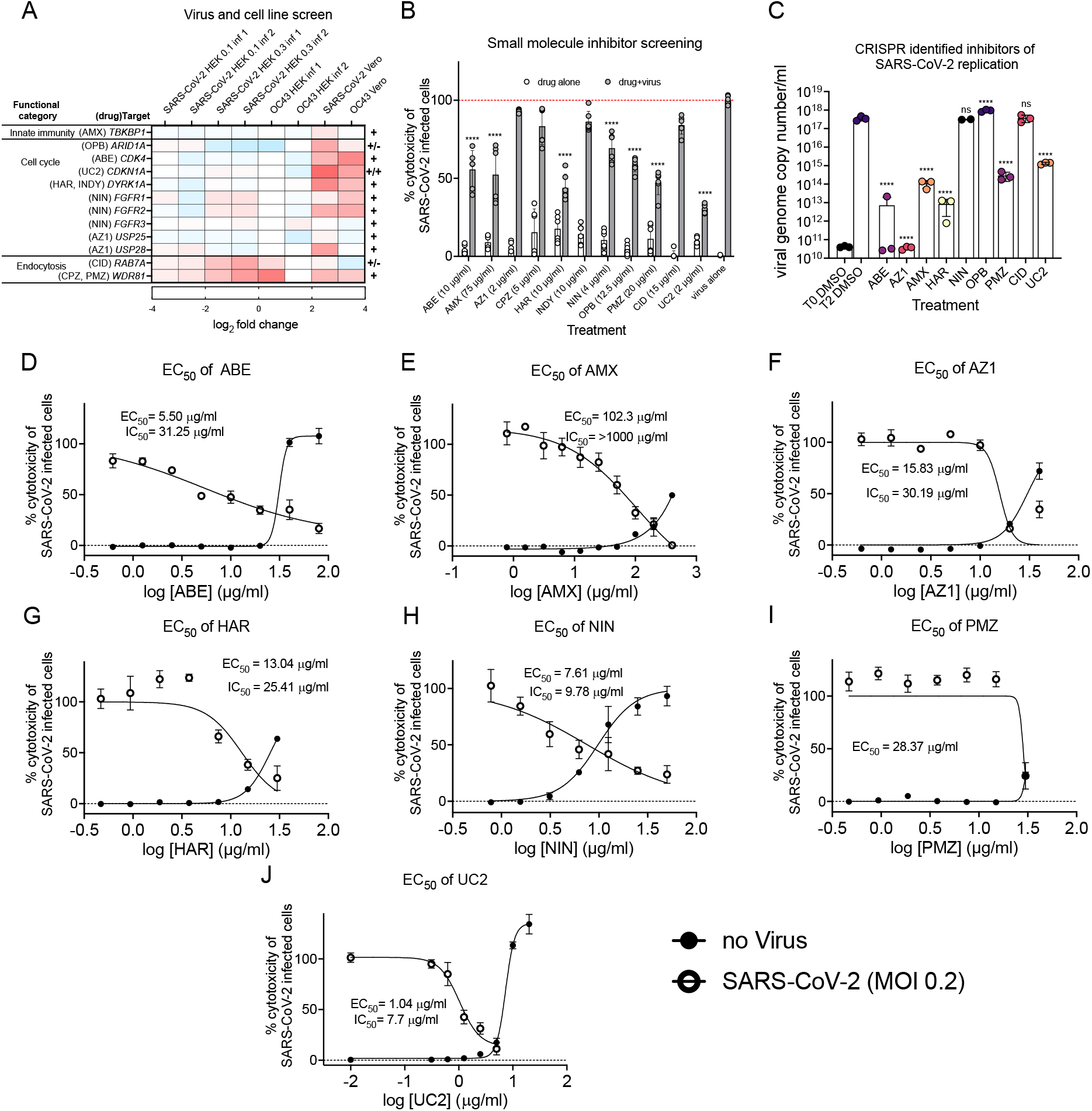
Small molecules to CRISPR-identified targets inhibit SARS-CoV-2 infection. **A)** The log_2_FC across the studies performed in this work for the 12 gene targets that had commercially available inhibitors. The + and – signs to the right of the heat map summarize the ability of these small molecules to prevent SARS-CoV-2 infection of Vero E6 cells. **B)** Initial screening of drug inhibition of SARS-CoV-2-induced cytotoxicity in Vero E6 cells following MOI 0.3 infection. **C)** Drug inhibition of SARS-CoV-2 genome replication was measured by RT-qPCR at 2 dpi of Vero E6 cells at MOI 0.01. One-way ANOVA was used to compare inhibitor-treated toxicity values to virus-alone controls. **D-J)** The EC_50_ (ability of inhibitors to reduce SARS-CoV-2 cytotoxicity) and IC_50_ (toxicity of inhibitors alone) curves were obtained by cytotoxicity assays in Vero E6 cells. Non-linear regression of the data points was used to determine the EC_50_ and IC_50_ values. Error bars indicate standard deviation for all panels (*n* = 3).

In an initial experiment of the entire panel of small molecules, inhibitors were added to culture supernatants at the initiation of SARS-CoV-2 infection and evaluated for their capacity to inhibit virus-induced CPE at 3 dpi in Vero E6 cells. The concentrations of inhibitors used, based on available toxicity data, were generally nontoxic in Vero E6 cells **(Fig. 7B, white bars).** ABE, AMX, HAR, NIN, OPB, PMZ, and UC2 significantly inhibited virus-induced cytotoxicity while AZ1, CPZ, INDY, and CID did not **(Fig. 7B, gray bars)**. Our CID results are consistent with prior work which showed reduced CoV egress, but no effect on cell viability or viral replication, in response to CID treatment *(2)*. Although INDY and HAR both target Dyrk1A, only HAR displayed activity in this assay, potentially due to the lower enzymatic IC_50_ of HAR for Dyrk1A (0.24 μM for INDY vs. 0.08 μM for HAR). As a complementary approach to measure antiviral activity of these compounds, we quantified viral genome copies by RT-qPCR in cells treated with each compound at 2 dpi. ABE, AMX, HAR, PMZ and UC2 significantly decreased viral genome copy number **(Fig. 7C)**, consistent with their ability to protect from virus-induced cytotoxicity. On the other hand, NIN and OPB had no effect on viral genome copy number despite their moderate inhibition of SARS-CoV-2-induced cytotoxicity. Conversely, AZ1 completely inhibited viral genome replication in spite of having no significant effect on cytotoxicity.

For compounds displaying activity in one or both of these assays, we next determined their EC_50_ and IC_50_ against SARS-CoV-2 infection by cytotoxicity measurements in the presence or absence of virus across a series of inhibitor dilutions **(Figs. 7D-J)**. Several of the inhibitors had EC_50_ values below 20 μM (10.86 μM for ABE, 14.1 μM for NIN, and 2.16 μM for UC2), with the p21 inhibitor UC2 being the most potent. AMX is typically used as topical treatment and had a high EC_50_ at 342.96 μM. AZ1 (37.49 μM), HAR (61.44 μM) and PMZ (88.41 μM) showed intermediate EC_50_ levels. Overall, these findings reveal novel candidates for anti-HCoV drug development.

## DISCUSSION

Genome-wide CRISPR knockout screens have been very successful in identifying host factors required for viral infection so it is not surprising that this approach has been applied to the discovery of proviral factors for SARS-CoV-2 infection. Indeed, six recent studies have reported CRISPR screens in cells infected with SARS-CoV-2 *(5–10).* As has been observed generally for genome-wide screens, there was limited overlap in the set of genes reported as host factors. In our reanalysis of the data using a common framework, there were only 17 genes identified in two or more published screens performed in human cells (*ACE2, ATP6V1A, ATP6V1G1, B3GAT3, CCZ1B, CNOT4, CTSL, DNM2, EXOC2, EXT1, EXTL3, MBTPS2, PIK3C3, SCAP, TMEM106B, VAC14*, and *VPS35*). This finding is not unexpected considering that the screens were performed in a variety of cell lines and under varying infection conditions. There was more overlap in the functional categories identified across studies, with enrichment of genes involved in glycosaminoglycan biosynthesis, vesicle transport, and ER/Golgi-localized proteins *(18).* Due to the limited redundancy of specific host factors identified across published studies and the potential of proviral gene products to be targeted with antiviral drugs, additional genome-wide screens are warranted. To that end, we performed CRISPR screens in AGM Vero E6 cells and human HEK293T-hACE2 cells. We performed screens for both SARS-CoV-2 and the common cold-causing HCoV OC43 to increase the probability of identifying pan-HCoV proviral factors representing strong targets for developing broad-spectrum antivirals. Our study provides additional support for previously identified candidate host factors and reports multiple novel host factors and pathways playing potentially key roles in viral replication. We summarize the consolidated set of candidate host factors identified for SARS-CoV-2 in our study as well as those identified in two or more studies in **Fig. 8**.

**Fig. 8.**
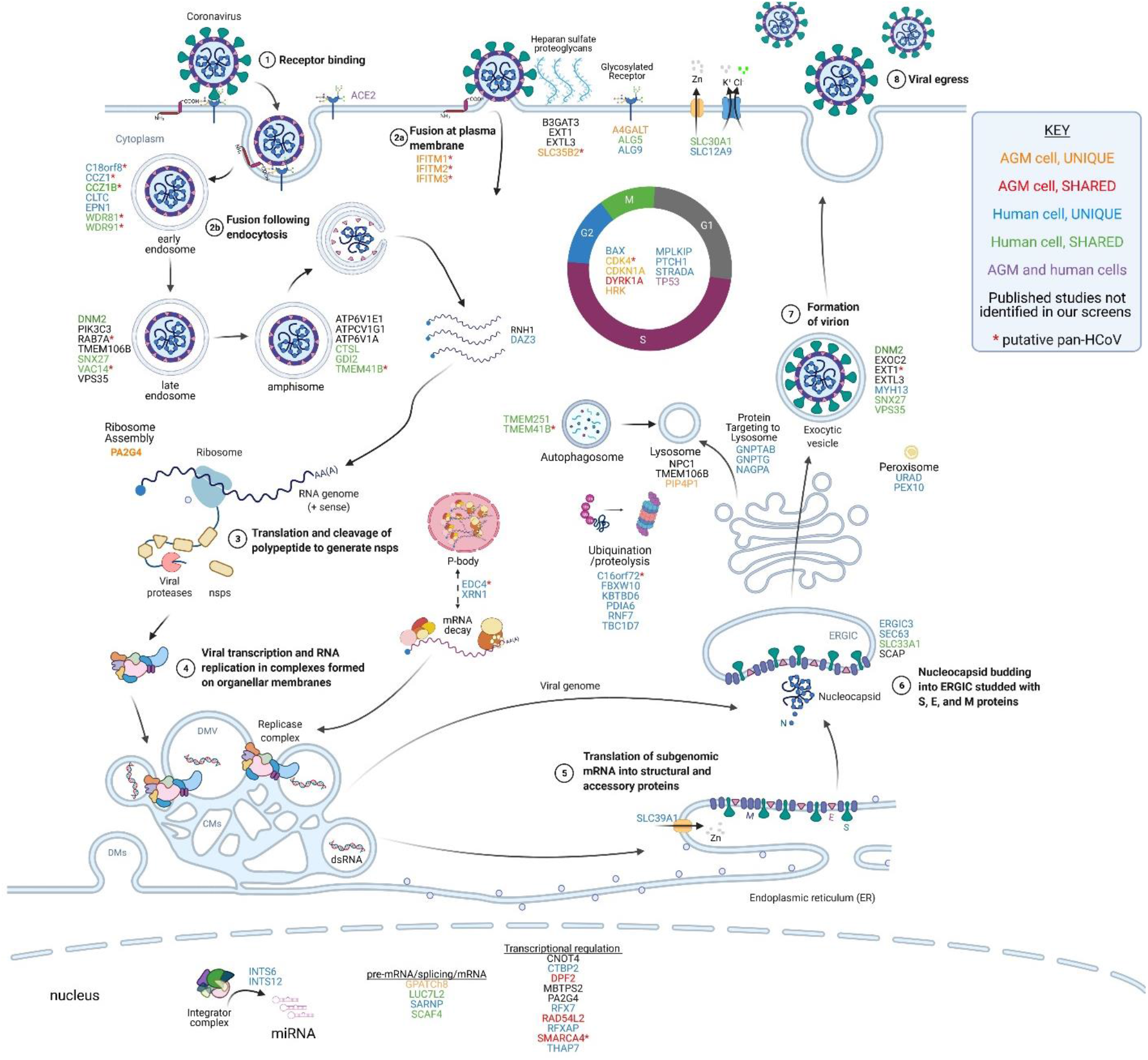
Summary of genes found in this and other studies and their potential roles in the SARS-CoV-2 life cycle. The host factors identified in CRISPR screens are presented adjacent to the putative stage of viral replication where they function. The genes are color-coded based on their identification in our and other published studies, as indicated in the legend. Candidate pan-HCoV host factors are indicated with red asterisks. The virus replicates through a series of well-defined molecular steps. **1-2)** After virion binding to ACE2, SARS-CoV-2 can fuse at the plasma membrane or following endocytosis. Heparan sulfate proteoglycans enhance viral attachment to cells so host factors involved in heparan sulfate biosynthesis (B3GAT3, EXT1, EXTL3, SLC35B2) and glycosylation (A4GALT, ALG5, ALG9) may play a role in viral entry. The IFITM proteins are proposed to promote fusion at the cell surface but inhibit fusion in endosomes. Host factors involved in endocytosis (C18orf8, CCZ1, CCZ1B, CLTC, EPN1, WDR81, WDR91), vesicular transport (DNM2, PIK3C3, RAB7A, TMEM106B, SNX27, VAC14, VPS35), and amphisome maturation/lysosome fusion (ATP6VIE1, ATPCV1G1, ATP6V1A, CTSL, GDI2, TMEM41B) likely facilitate virion uncoating. **3)** The positive-sense RNA genome is then translated to produce the nonstructural polyproteins which are co-translationally cleaved to form the mature nsps. Certain host factors like RNH1 and DAZ3 may serve to protect the viral genome from degradation by host enzymes. **4)** The nsps form the viral replicase which assembles on organellar membranes to form the replication and transcription complexes (RTCs) where progeny genomes and structural/accessory protein transcripts are produced, respectively. P-body components EDC4 and XRN1, identified in this study, may play a role in the maintaining viral RNA stability or assembly of the RTC. **5)** Structural and accessory proteins are translated, and structural proteins insert into the ER membrane. ER-localized SLC39A1 may play a role in this process. **6)** Nucleocapsids bud into the ERGIC, potentially aided by host factors ERGIC3, SEC63, SLC33A1, and SCAP. **7)** Progeny virions form as they traverse through the Golgi and structural proteins are glycosylated. **8)** Virions exit the cell through either typical exocytosis (DNM2, EXOC2, EXT1, EXTL3, MYH13, SNX27, VPS35) or nonclassical lysosomal egress (GNPTAB, GNPTG, NAGPA, NPC1, TMEM106B, PIP4P1). Numerous host factors with less obvious direct roles in promoting steps in the viral life cycle have also been identified in CRISPR screens. For example, numerous factors regulating the cell cycle (BAX, CDK4, CDKN1A, DYRK1A, HRK, MPLKIP, PTCH1, STRADA, TP53) were identified in our screens in AGM and human cells. Furthermore, multiple nuclear-localized host factors including diverse transcriptional regulators and two components of the integrator complex (INTS6, INTS12) were identified. Overall, the large number of diverse host factors that promote SARS-CoV-2 replication illustrates the large-scale exploitation of cellular processes required for successful viral propagation. Adapted from BioRender template titled Life Cycle of Coronavirus generated by the Britt Glaunsinger laboratory. Created with BioRender.com.

Host factors promoting SARS-CoV-2 infection of Vero E6 cells (FDR<0.25) are indicated in **Fig. 8** adjacent to the presumptive step in the viral life cycle in which they function. In this cell line, cell cycle regulation was key to SARS-CoV-2 replication. *CDK4* was a top-scoring gene for both SARS-CoV-2 and OC43, suggesting it is broadly required for HCoV replication. CoVs utilize diverse strategies to manipulate the host cell cycle to promote their replication *(21)*. Identification of specific cell cycle-related host factors required for HCoV replication could provide clues to dissecting viral regulatory mechanisms. We also identified IFITM proteins in both SARS-CoV-2 and OC43 screens in Vero E6 cells, consistent with a prior study reporting that IFITM proteins promote OC43 infection *(12)*. Interestingly, recent work suggests that IFITM proteins promote HCoV entry when it occurs at the plasma membrane but inhibit HCoV entry when it occurs in the endocytic pathway *(13, 14)*, suggesting that HCoVs enter Vero E6 cells primarily at the plasma membrane instead of using the endosomal pathway. This finding is consistent with the paucity of factors involved in endocytosis identified in these screens, in stark contrast to our and others’ results in screens performed in human cell lines.

In addition to CDK4 and IFITM proteins, targeting of *SLC35B2* in both SARS-CoV-2 and OC43 Vero E6 screens increased resistance to infection, suggesting that it is a pan-HCoV host factor in this cell line. *SLC35B2* encodes 3’-phosphoadenosine 5’-phosphosulfate transporter 1 (PAPST1) which plays an important role in heparan sulfate biosynthesis. PAPST1 is required for optimal replication of a variety of viruses including HIV, dengue virus, and bunyaviruses, enabling heparan sulfate-mediated viral entry or sulfating a viral receptor that enables virion binding *(22–24)*. It is hence logical to predict that it functions in HCoV entry in Vero E6 cells as well. Additional candidate pan-HCoV factors identified in the Vero E6 studies include *PLN* encoding phospholamban and *C16orf74* which are both implicated in maintaining calcium homeostasis *(25–27)*, and *C3orf80* encoding a protein of unknown function. None of these gene products have been previously identified as viral host factors to our knowledge, and their functional roles will require further study.

Host factors promoting SARS-CoV-2 infection of HEK293T-hACE2 cells (FDR<0.25) are indicated in **Fig. 8** adjacent to the presumptive step in the viral life cycle in which they function. The functional categories with the most top-scoring genes were vesicle transport, cell cycle regulation, autophagy, and ubiquitination/proteolysis. For the OC43 screens, the most abundant functional categories were vesicle transport, transcriptional regulation including the SWI/SNF complex, innate immunity, and transporters. The host factors identified in the HEK293T-hACE2 screens for both SARS-CoV-2 and OC43 (C18orf8, CCZ1, CCZ1B, RAB7A, WDR81, and WDR91) are all involved in vesicle-mediated transport and particularly in endosomal maturation, underscoring the importance of this process for HCoV infection.

When comparing our SARS-CoV-2 data sets in HEK293T-hACE2 cells to published data sets in other human cell lines, there were 21 genes in common with other studies (FDR<0.25; *ACE2, ALG5, ARVCF, CCZ1B, CTSL, DNM2, EPT1, GDI2, LUC7L2, RAB7A, RNH1, SCAF4, SLC30A1, SLC33A1, SNX27, TMEM41B, TMEM251, VAC14, VPS35, WDR81*, and *WDR91*) which highlight key functional pathways required for viral infection, including endocytosis, glycosylation, and exocytosis. Remarkably though, we identified 53 unique genes, underscoring the importance of continued screening to fully elucidate host factors promoting SARS-CoV-2 replication. Certain unique genes function in previously identified pathways such as vesicle transport (e.g., *CCZ1, C18orf8*) and ER/Golgi-localized proteins (e.g., *SEC63, ERGIC3*). Other unique genes function in processes that have not been previously described as proviral in HCoV infections. For example, *EDC4* was a top-scoring gene in our SARS-CoV-2 screens in HEK293T-hACE2 cells. EDC4 functions as a scaffold protein for the assembly of the programmed mRNA decay complex. Although it has not been reported to play a role in HCoV infection before, it does promote rotavirus replication complex assembly *(28)*. Another component of the programmed mRNA decay pathway XRN1 was also modestly enriched, suggesting that this pathway promotes SARS-CoV-2 replication. Alternatively, EDC4 and XRN1 are both P-body components. Many RNA viruses interact with and hijack P-bodies in order to promote viral replication *(29)* and SARS-CoV-2 has recently been reported to disrupt P-bodies *(30)* so it is possible that the virus interacts with these host factors to disassemble P-bodies and facilitate viral replication. We also identified three unique genes encoding factors involved in targeting proteins to lysosomes – GNPTAB, GNPTG, and NAGPA. Considering recent progress in understanding the key role played by lysosomes in HCoV egress *(2)*, it is interesting to speculate that HCoVs interact with these proteins to facilitate virion release from the infected cell.

Two approaches were taken to validate the proviral role of a subset of unique host factors identified in our screens. First, shRNA-mediated knockdown of *CCZ1* and *EDC4* resulted in reduced SARS-CoV-2 and OC43 replication. Second, drugs targeting selected host factors displayed antiviral efficacy *in vitro* against SARS-CoV-2. These include cell cycle inhibitors ABE targeting Cdk4, AZ1 targeting Usp25/28, HAR targeting Dyrk1A, NIN targeting Fgfr1/2/3, and UC2 targeting p21; the endocytosis inhibitor PMZ targeting Wdr81; and the Tbk1 inhibitor AMX. Chen et al. recently reported similar activity of ABE against SARS-CoV-2 *(31),* validating our findings. To our knowledge, the discovery that AMX, AZ1, HAR, NIN, PMZ, and UC2 possess antiviral activity against SARS-CoV-2 has not been reported. AMX, PMZ, and NIN are currently available drugs which could potentially be repurposed, while HAR is a natural product being investigated for the treatment of a variety of diseases. While no clinical therapeutics are currently available targeting p21 or Usp25/28, our data suggest that these could be worthwhile targets for drug development. Further study of the potential *in vivo* utility of these compounds in treating HCoV infections and their mechanism of action is warranted.

Our studies substantiate and expand the growing body of literature focused on understanding key HCoV-host cell interactions. The fairly limited redundancy in proviral factors identified across our study and other published studies using genome-wide CRISPR screens *(5–10)* highlights the extensive scope of these interactions and suggests that even more host factors remain to be discovered. Cell type differences and variable infection conditions undoubtedly influence the outcomes of screens and could provide novel insight into nuanced viral replication mechanisms. For example, we identified lysosomal proteins as proviral in HEK293T-hACE2 cells but not in Vero E6 cells, raising the possibility that there are cell type-specific differences in the use of lysosomes for HCoV egress *(2)*. Detailed molecular studies testing hypotheses like this stemming from genome-scale CRISPR screens are a critical next step. Similarly, although we have identified novel compounds displaying antiviral activity against HCoVs *in vitro*, additional work is needed to determine their mechanism of action at the molecular level and *in vivo* efficacy before they can be applied in the clinic.

## MATERIALS AND METHODS

### Study design

The objectives of this study were to identify host factors that promote HCoV replication and to determine whether these host factors can be targeted with commercially available drugs to block viral infection *in vitro* **(Fig. S1)**. To achieve this goal, we performed genome-wide CRISPR knockout screens in Vero E6 cells using a newly generated Vervet sgRNA library **(Fig. 1A)** and in HEK293T-hACE2 cells using the commercially available human Brunello sgRNA library **(Fig. 1B)**. In brief, cells transduced with sgRNA libraries were infected with HCoVs, SARS-CoV-2 and OC43, using various MOIs. Cells surviving infection were expanded and re-infected, and the sgRNAs enriched in resistant clones was determined. Genes targeted by enriched sgRNAs were compared between replicates, infection conditions, HCoVs, cell lines, and previously published screens to identify common and unique host factors as well as putative pan-HCoV host factors. Genes of interest were selected for validation and targeted for knockdown using shRNAs followed by infection with HCoVs. Commercially available inhibitors to other important genes identified in our study were evaluated for their capacity to prevent virus-induced toxicity and viral replication *in vitro*. EC_50_ and IC_50_ values for efficacious compounds were determined.

### Virus stock generation and titer determination

SARS-CoV-2 strain UF-1 (GenBank accession number MT295464.1) was originally isolated from a COVID-19 patient at the University of Florida Health Shands Hospital via nasal swab *(32)* and manipulated in a Biosafety Level 3 (BSL3) laboratory at the Emerging Pathogens Institute under a protocol approved by the University of Florida Institutional Biosafety Committee. The HCoV OC43 strain was a kind gift from Dr. John Lednicky (University of Florida). SARS-CoV-2 and OC43 were propagated in Vero E6 cells (ATCC) grown in Dulbecco’s Modified Eagle Medium (DMEM; Gibco) supplemented with 10% heat inactivated fetal bovine serum (FBS; Atlanta Biologicals) and Pen-Strep (100 U/ml penicillin, 100 μg/ml streptomycin; Gibco) at 37°C and 5% CO_2_. Virus stocks were prepared by infecting Vero E6 cells at MOI 0.01, centrifuging culture supernatants collected at 3 dpi for 5 mins at 1000 x g, and filtering through a 0.44 μm PVDF filter (Millipore) followed by a 0.22 μm PVDF filter (Restek). The virus stocks were aliquoted and stored at −80°C. Virus stocks were titered using a standard TCID50 assay. In brief, Vero E6 cells were seeded at 2×10^4^ cells per well in a 96-well plate (Corning) and allowed to attach overnight. Virus stocks were serially diluted onto cells, with a total of 8 replicates per dilution. Monolayers were visualized in the BSL3 using an EVOS XL Core microscope (Thermo Fisher Scientific) and scored positive or negative for cytopathic effect (CPE) at 7 dpi.

### Viral genome copy number enumeration

For SARS-CoV-2, supernatants and cells were harvested into AVL buffer from the QIAamp Viral RNA Kit (Qiagen) and RNA was purified according to the manufacturer’s recommendations. The samples underwent reverse transcription and cDNA synthesis using the iTaq Universal SYBR Green One-Step Kit (BioRad) and primers targeting the nucleocapsid (N) gene of SARS-CoV-2 (NproteinF-GCCTCTTCTCGTTCCTCATCAC, NproteinR-AGCAGCATCACCGCCATTG). qPCR was carried out on a Bio-Rad CFX96 and viral genome copy numbers were extrapolated using CT values from a standard curve generated using a control plasmid containing the N protein gene (Integrated DNA Technologies). For OC43, RNA from infected cells was purified using the RNeasy Mini Kit (Qiagen) according to the manufacturer’s recommendations and amplified using the Applied Biosystems AgPath-ID One-Step RT-PCR Kit (Thermo Fisher Scientific) and primers and probe targeting the N gene of OC43 *(33).* GAPDH levels were determined for each sample for normalization purposes using previously described primers *(34)*. All samples were run in triplicate for each primer pair and normalized viral genome copy numbers were calculated using the comparative cycle threshold method.

### Genome-wide CRISPR sgRNA screens

The human CRISPR Brunello library (Addgene 73178) *(16)* was amplified following a previously published protocol *(35)*. We constructed a Vervet domain-targeted sgRNA library since one was not commercially available. Detailed methods are reported in supplemental material. For both libraries, lentiviruses were produced in HEK293T cells by co-transfection of library plasmids together with the packaging plasmid psPAX2 (Addgene 12260) and envelope plasmid pMD2.G (Addgene 12259). CRISPR screens were carried out in two cell lines (outlined in **Fig. 1**): AGM Vero E6 cells were transduced with the newly generated Vervet sgRNA library and human HEK293T-hACE2 cells (Genecopoeia) were transduced with the Brunello sgRNA library. For each screen, 1.2×10^8^ cells were transduced with lentivirus-packaged sgRNA library at MOI 0.3 in the presence of 8 μg/ml polybrene (Sigma) to achieve ~500-fold overrepresentation of each sgRNA. After 48 h, 0.6 μg/ml puromycin (Gibco) was added to eliminate non-transduced cells and cultures were expanded in Matrigel-coated (Corning) T300 flasks. Control replicates were collected at this time to determine input library composition and additional replicates were infected with SARS-CoV-2 or OC43 at the indicated MOIs. Cells surviving initial infections were collected when they had expanded to confluency. A portion of each replicate was stored in DNA/RNA Shield (Zymo Research) at −80°C for genomic DNA extraction and the remaining cells were reseeded and reinfected at the indicated MOIs. Cells surviving reinfections were also harvested at confluency for genomic DNA extraction.

Genomic DNA was extracted from each sample (detailed extraction methods are described in supplemental methods). The sgRNA regions were then amplified and indexed for Illumina sequencing using a one-step PCR method and primers specific to the LentiCRISPRv2-based Vervet and Brunello libraries. Primers and indices used for the generation of amplicon libraries are listed in **Table S1**. Brunello and Vervet DNA samples were amplified in ten 100 μl reactions using the NEBNext High-Fidelity 2X Master Mix Kit (New England Biolabs), 0.5 μM of forward and reverse primers and 10 μg of DNA template per reaction with the following program: initial denaturation at 98°C for 3 min, 24 cycles of denaturation at 98°C for 10 s, annealing 60°C for 15 s and extension 72°C for 25 s, and final extension at 72°C for 2 min. 256 bp amplicons were quantified on a 2% agarose gel stained with SYBR Safe (Invitrogen), using the Gel Doc quantification software (Bio-Rad). Amplicons were first pooled in an equimolar fashion and then the pools were gel-extracted using the PureLink Quick Gel Extraction Kit (Thermo Fisher Scientific). The sequencing was carried out at the Interdisciplinary Center for Biotechnology Research (ICBR; University of Florida) using a NovaSeq 6000 sequencer (Illumina). The Brunello amplicons were sequenced using the S4 2X150 cycles Kit (Illumina) while the Vervet-AGM amplicons were sequenced with the SP 1X100 cycles Kit (Illumina).

### Computational analysis

The FASTX-Toolkit was used to demultiplex raw FASTQ data which were further processed to generate reads containing only the unique 20 bp sgRNA sequences. The sgRNA sequences from the library were assembled into a Burrows-Wheeler index using the Bowtie build-index function and reads were aligned to the index. The efficiency of alignment was checked and the number of uniquely aligned reads for each library sequence was calculated to create a table of raw counts. Ranking of genes corresponding to perturbations that are enriched in infected cultures was performed using a robust ranking aggregation (a-RRA) algorithm implemented in the Model-based Analysis of Genome-wide CRISPR/Cas9 Knockout (MAGeCK) tool through the test module. Tables with raw counts corresponding to each sgRNA in reference (initial pool) and selected (virus-infected) samples were used as an input for MAGeCK test. Gene-level ranking was based on FDRs and candidates with FDRs < 0.25 were considered as significant hits. Ranking of genes corresponding to positively selected and negatively selected perturbations was performed using a robust ranking aggregation (a-RRA) algorithm implemented in MAGeCK through the test module *(36)*. Tables with raw counts corresponding to each sgRNA in reference (initial pool) and selected (exposed to virus) samples were used as an input for MAGeCK test. Gene-level ranking was based on false discovery rate (FDR) and candidates with FDR < 0.25 were considered as significant hits. Additional details can be found in the supplementary methods. We submitted FastQ files to Gene expression omnibus (GSE: XXXXXX) and all CRISPR screen data to BioGRID ORCS database (https://orcs.thebiogrid.org/). In addition, all data is available at sarscrisprscreens.epi.ufl.edu.

### Validation of host factors in promoting viral infections

To validate selected host factors for their capacity to promote HCoV infection *in vitro*, we transduced HEK293T-hACE2 cells with lentivirus-packaged shRNAs targeting *CTSL, CCZ1*, or *EDC4* (TRC Human shRNA Library collection) or the empty vector pLKO.1. Transduced cells were expanded under puromycin selection (0.8 ug/ml) for at least 7 days to generate stable knockdown cell lines. To confirm knockdown, cell lysates prepared from knockdown and control cells were tested by western blotting with antibodies directed to CTSL (Invitrogen, BMS1032), CCZ1 (Santa Cruz Biotechnology, sc-514290), EDC4 (Cell Signaling Technology, 2548S), and actin as a loading control (Sigma-Aldrich, MAB1501R). Once knockdown was confirmed, cells were infected with SARS-CoV-2 or OC43 at MOI 0.01 and RNA was extracted at 2 dpi for viral genome copy number enumeration, as described above.

### Identification and testing inhibitors of host factors from CRISPR screens

Online databases and published literature were used to find small molecule inhibitors targeting a subset of top-scoring genes in the CRISPR screens. Amlexanox was purchased from InvivoGen. Abemaciclib, AZ1, carfilzomib, olaparib, and nintedanib were purchased from Selleckchem. Harmine, INDY, chlorpromazine, promethazine, UC2288, and CID 1067700 were purchased from Millipore Sigma. Drugs were diluted according to the manufacturers’ recommendations and single-use aliquots were frozen at −80°C until the time of assay. Drugs were diluted down to 2X concentrations and mixed with 2X concentrations of virus to generate 1X concentrations, then added to the monolayers. Cells were infected at a MOI of 0.2 as determined by preliminary experiments to generate an ideal dynamic range of the colorimetric CytoTox 96 ® Non-Radioactive Cytotoxicity Assay (Promega). Infections progressed for 72 h at which time supernatant from treatments and controls were processed for LDH release according to the manufacturer’s recommendations. Absorbance at 450 nm was read using an accuSkan FC microplate reader (Fisher Scientific) with SkanIt software (Fisher Scientific). Absorbance values were background subtracted and transformed to percent of virus-infected controls. These percentages were compared to the values obtained from virus infected-cell cytotoxicity values by one-way ANOVA using GraphPad Prism version 9. Assays were carried out in biological duplicate and in three independent experiments. The concentration of drug alone that resulted in 50% of maximum toxicity (inhibitory concentration 50; IC_50_) and the concentration of drug that inhibited 50% of the vehicle treated SARS-CoV-2 induced cytotoxicity (effective concentration 50; EC_50_) were determined by serially diluting small molecule inhibitors during SARS-CoV-2 infection of Vero E6 cells. EC_50_ and IC_50_ values were calculated by transforming inhibitor concentrations to log then using the non-linear fit with variable slope function (GraphPad Prism version 9) to determine best fit variables using the percent of maximum SARS-CoV-2 induced cytotoxicity measurements at each drug concentration performed in technical duplicate.

### Website creation and data repository

To facilitate the access, reusability and integration of this data, we have created and hosted a website (sarscrisprscreens.epi.ufl.edu) which contains data for previously published HCoV CRISPR screens and our integrated MAGeCK analysis using the VISPR pipeline of the SARS-CoV-2 screens. Our goal is to provide a community resource for facile integrated analysis of current and future CRISPR screens. Further details on how to submit new data is provided on the website and in supplemental methods.

### Statistical analysis

For the genome-scale CRISPR analysis, the embedded statistical tools in the MAGeCK/VISPR pipelines were used *(15, 36)*. Further details are provided in the supplemental materials. All other statistical analyses were carried out using GraphPad Prism 9.0. To compare the mean normalized viral genome copy number values in targeted shRNA knockdown experiments (Fig. 6), *P* values were determined using one-way ANOVA test (\**P* < 0.05, \*\**P* < 0.01, \*\*\**P* < 0.001), with error bars representing standard errors of mean (*n* = 3 experiments). For testing inhibitory activity of small molecules on SARS-CoV-2 infection of Vero E6 cells (Fig. 7), a one-way ANOVA test was used for comparison of toxicity values for inhibitor-treated infected cells and infected-only control cells (no treatment), with error bars denoting standard deviation for all panels (*n* = 3 experiments). Non-linear regression of data points was used to determine the EC_50_ and IC_50_ values for indicated compounds.

## Acknowledgments

We thank Dr. John Lednicky (University of Florida, UF) and the UF Health Cancer Center for providing key reagents. This work was supported by the UF Clinical and Translational Science Institute. S.M.K. and M.G. were further supported by NIH R01AI123144.

## Author contributions

Conceptualization: MHN, SMK, CDV

Methodology: MG, APB, MS, AT, MHN, SMK, CDV

Investigation: MG, APB, MS, AT, MR, AS, RR, MHN

Funding acquisition: MHN, SMK, CDV

Project administration: MHN, SMK, CDV

Supervision: MHN, SMK, CDV

Writing – original draft: MG, APB

Writing – review & editing: MHN, SMK, CDV

## Data and materials availability

All data are available in the main text or the supplementary materials.

